# Canonical Wnt/β-catenin signalling regulates inducible aging and regeneration loss in hydra

**DOI:** 10.1101/2025.04.01.646560

**Authors:** Jácint Tökölyi, Yukino Kumagai, Regina Krisztina Szilágyi, Nikoletta Andrea Nagy, Virág Szakál, Retno Novvitasari Hery Daryono

## Abstract

Freshwater cnidarians from the genus *Hydra* have exceptional regeneration capacities and show negligible aging. However, one species, *Hydra oligactis*, experiences accelerated senescence following sexual reproduction, characterized by regeneration loss, stem cell depletion, reduced body size, motility and food capture rates. This phenomenon, termed inducible aging, is triggered by temperature-induced sexual reproduction. The physiological regulation of the switch from high regenerative capacity and low senescence to regeneration loss and accelerated aging remain largely unexplored. By comparing gene expression patterns of asexual and sexual polyps following amputation we identified several canonical Wnt/β-catenin signalling pathway transcripts that showed differential expression in the regeneration-deficient sexual individuals, suggesting the involvement of this pathway in the inducible aging phenotype. Pharmacological activation of canonical Wnt/β-catenin signalling with alsterpaullone (ALP) restored head regeneration and improved survival of animals. To find out more about the role of this pathway in sexual development and post-reproductive senescence, we treated animals in various stages of egg maturation with ALP. We found that ALP delayed egg maturation when applied in early stages, but had smaller effects when applied in later stages of gametogenesis, without having a stark effect on overall fecundity. ALP treatment increased survival following sexual reproduction. These results show that the canonical Wnt/β-catenin signalling pathway regulates reproduction, regeneration and post-reproductive senescence in *H. oligactis*.

## Introduction

Animals display extraordinary variation in their patterns of growth, reproduction and survival - collectively known as life history traits. Despite their remarkable diversity, life history traits do not vary freely but appear intrinsically linked. For instance, reduced growth and reproductive rates often correlate with extended longevity^1,2^, while cases of sustained reproductive effort alongside extended longevity are rare^3^. Such correlations have led to the view that life history traits are not regulated by separate genetic mechanisms, but instead result from pleiotropic physiological regulatory pathways that balance complementary investments into survival and reproduction to maximize Darwinian fitness under specific ecological conditions^4–6^.

Examples of pleiotropic regulatory mechanisms have been detected in organisms ranging from plants and algae^7,8^, through invertebrates^9–11^ to mammals^12–14^. Most prominently, the insulin/insulin-like signalling pathway (IIS) and the mechanistic Target of Rapamycin (mTOR) have been implicated in regulating cell proliferation, growth, reproductive investment and longevity in multiple animal groups^10,14–16^. For instance in flies, mutations in the IIS pathway result in juvenile hormone deficiency, which can retard growth and result in extended longevity^17^. Likewise, autophagy regulated by the mTOR pathway is critical for normal development early in life in mice, but the same pathway contributes to senescence later in life^16^. Recently, canonical Wnt signalling has been implicated in regulating the trade-off between egg-laying, asexual reproduction and regeneration in flatworms^18^.

While these pleiotropic regulatory mechanisms have received substantial attention in model organisms, the diversity of physiological mechanisms regulating fitness traits and the longevity-reproduction trade-off - both in terms of phylogenetic diversity and at the level of the mechanisms involved - is still to be deciphered in detail. The freshwater cnidarian hydra is one of the few unusual animal species that has been shown to exhibit extended longevity while also maintaining high reproduction ^3,19,20^. *Hydra vulgaris* (Pallas, 1766) polyps have been followed for nearly a decade in the laboratory, during which they experienced negligible mortality despite maintaining a constant rate of asexual reproduction. Based on their patterns of mortality, the expected maximum lifetime potential of *Hydra vulgaris* was estimated to be >3000 years^19^ and is considered to be the result of high self-renewal capacity of hydra polyps due to a high proportion of stem cells and excessive regeneration capacity^19,21^.

Interestingly, another species from the same genus, *Hydra oligactis* (Pallas, 1766), also assumed to have extended longevity when reproducing asexually^22^, experiences increased mortality following sexual reproduction, a process induced by a sudden drop in temperature (Fig. 1A). This phenomenon, termed inducible aging, is associated with stem cell depletion, loss of regeneration ability, reduced motility and food capture, and ultimately, a high probability of death just within a few months after initiation of sexual reproduction^22–25^. However, some individuals survive gametogenesis and revert to asexual reproduction while still maintained at low temperatures^26^. Hence, senescing and non-senescing individuals from the same genotype can be studied under identical environmental conditions, which makes this an ideal model system to investigate the physiological regulatory mechanisms behind stem cell differentiation patterns, regeneration loss and aging.

**Fig. 1.**
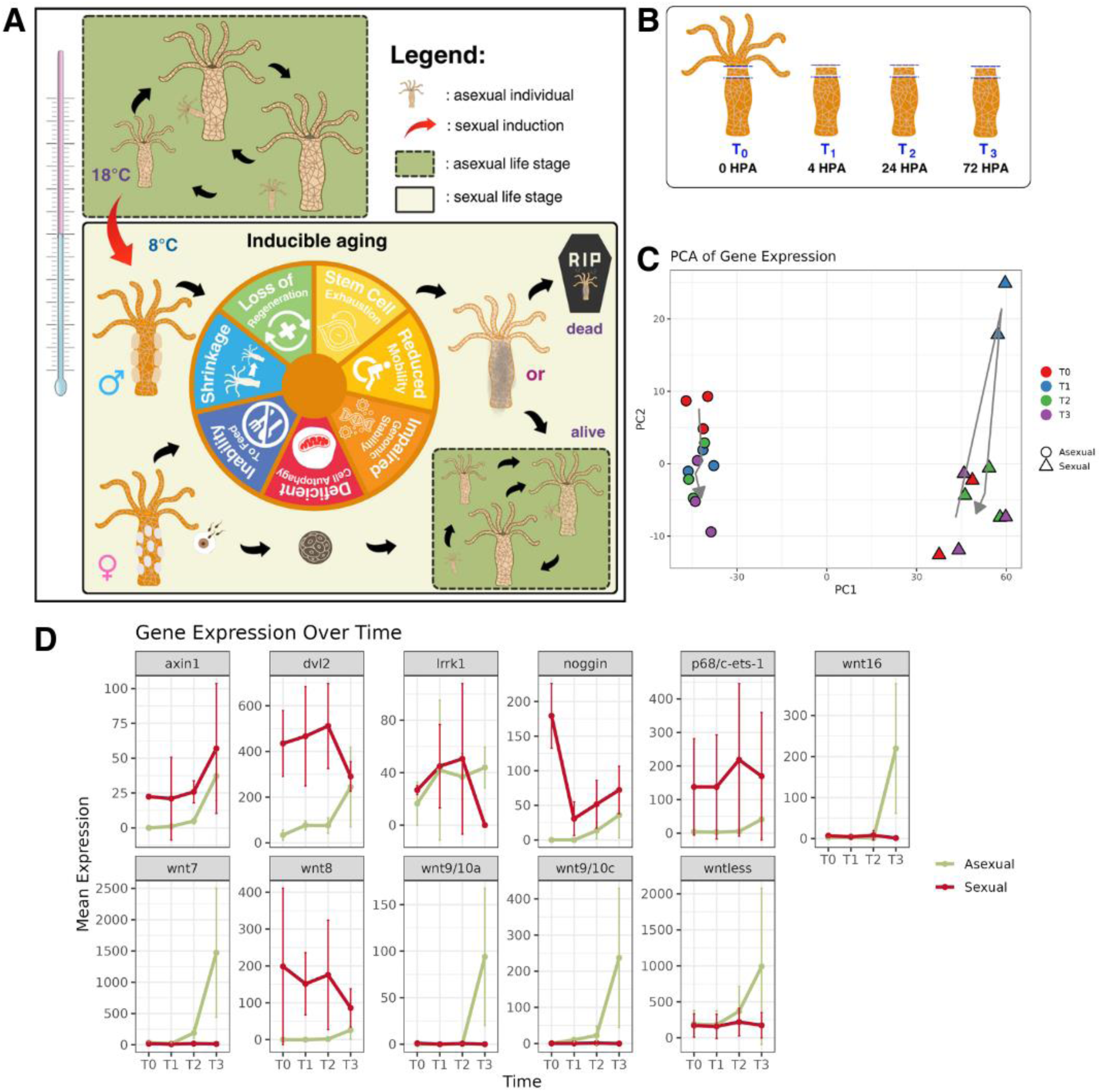
Transcriptomic changes in response to head amputation in asexual (regenerating) and sexual (regeneration-deficient) hydra. A) Schematic showing the sexual and asexual life stages of Hydra oligactis. Polyps reproduce asexually at higher temperatures through budding. Lowering the temperature results in a switch from asexual to sexual reproduction and egg production in around three weeks. Sexual reproduction is associated with reduction in body size, regeneration loss, stem cell depletion, inability to feed and move. Some of the polyps will die, but others can recover and revert to asexual reproduction at low temperature, while maintaining high regeneration ability. B) Transcriptomic changes following head amputation were investigated in the tissue below the amputation plane at 0, 4, 24 and 72 hpa, in both asexual (regenerating) and asexual (regeneration-deficient), both cultured at 8 °C. C) PCA showing overall gene expression differences. D) expression pattern of canonical wnt/β-catenin signalling pathway genes that showed significant Time * Reproductive mode interaction in their expression pattern.

Previous studies have implicated loss of autophagy^27^, loss of genome maintenance^25^ and stem cell depletion^23,25,28^ behind inducible aging in *Hydra oligactis*. However, identifying the underlying mechanism proved difficult due to complex differences in the physiology of sexual (aging) and asexual (non-aging) forms, lack of tools to interrogate the genetics of the species and gaps in the knowledge of its life cycle. To address this problem, we studied gene expression patterns of sexual (non-regenerating) and asexual (regenerating) hydra polyps from the same strain following amputation. We hypothesized that altered function of canonical Wnt/β-catenin signalling, a central regulator of cell differentiation and regeneration in hydra^21,29,30^, might be the mechanism behind regeneration loss in aging hydra. Furthermore, we used pharmacological activation of Wnt signalling, to understand how the canonical Wnt/β-catenin pathway regulates the inducible aging phenotype in hydra.

## Methods

### Hydra strain & housing

For all experiments reported here, we used the female strain X11/14. This strain was established from a polyp collected in Hungary in 2016 and maintained asexually since then in the lab. Animals were cultured individually in 6-well tissue culture plates in artificial hydra medium (1 mM Tris, 1 mM NaCl, 1 mM CaCl_2_, 0.1 mM KCl, 0.1 mM MgSO_4_; pH: 7.6) at 18 °C and 12/12 hours light/dark cycle. They were fed three times per week with freshly hatched *Artemia salina* nauplii. To induce sexual reproduction and obtain regeneration-deficient, aging individuals, they were transferred to an incubator set to 8 °C and an 8/16 hours light/dark cycle (sexual induction, SI). For some of the experiments reported here, we also used regeneration-capable asexual individuals cultured at 8 °C. These individuals belonged to strain X11/14 and were obtained from polyps surviving sexual reproduction in the lab and subsequently cultured under identical conditions as other polyps, except for the temperature difference (8 °C vs. 18 °C).

### RNA isolation from regenerating tissues

RNA samples were collected from sexual animals (2 weeks after transferring them from high to low temperature) and asexual animals. Both groups were cultured at 8 °C before and during the experiment. Regenerating tissue was collected at four distinct time points (Fig. 1B): 1) immediately after amputation (0 hours post-amputation, hpa); 2) 4 hours later (4 hpa); 3) 24 h later (24 hpa) and 4) 72 hours later (72 hpa). First, the head of all experimental animals was amputated with a sterile scalpel and discarded. Next, a ring of tissue corresponding to ∼10% of the body column just below the amputation plane was excised and immediately immersed in TrizolⓇ (Life Technologies, Carlsbad CA, USA). Tissue from four animals was used for one sample to increase RNA yield. RNA was extracted with TrizolⓇ following the protocol described in^31^ and subsequently sent to Novogene (Cambridge, UK) for 150 nt paired-end transcriptome sequencing. We aimed at having three biological replicates per time point in both the sexual and asexual groups (N=24 samples in total), however we failed to extract sufficient RNA from two samples (Sexual T0 and Sexual T1), making the final sample size N=22.

### Transcriptome assembly

Raw reads were subjected to several filtering steps to remove low-quality and contaminant sequences. First, we used *fastp v0.24.0* ^32^ to perform adapter detection, error correction, and trimming, while applying quality filtering criteria of a minimum Phred score of 30, requiring sequences to be at least 50 bases long, and trimming based on a sliding window of size 4 with a mean quality score of 30. Next, we used *SortMeRNA v4.3.7*^33^ to identify and remove rRNA contamination, by aligning sequences to the *smr_v4.3_sensitive* database. Finally, to identify sequences from other contamination sources (e.g. hydra’s microbiome, food or human tissue), we aligned reads to a high quality complete genome sequence of *H. oligactis*^34^ using *HISAT2 v2.2.1*^35^ and discarded non-aligning reads. The resulting set of quality-controlled, decontaminated reads was used for *de novo* transcriptome assembly with *Trinity v2.9.1*^36^, followed by transcript quantification using *salmon v1.10.0*^37^. From the assembled transcripts peptide sequences were identified with *TransDecoder v5.7.1* (https://github.com/TransDecoder/TransDecoder) and annotated with *Trinotate v4.0.2*^38^, by performing *blastp* and *blastn*^39^ searches in the UniprotKB/SwissProt database.

### Differential expression analysis

Differential gene expression analysis was conducted with the *DESeq2 v1.44.0* R package^40,41^ at the transcript (isoform) level. Transcripts with a total count across samples <10 and not present in at least 3 samples were filtered out. We initially compared overall gene expression at 0 hpa (T0), 4 hpa (T1), 24 hpa (T2) and 72 hpa (T3) in both sexual and asexual individuals using Principal Components Analysis. Subsequently, we identified genes differentially regulated in sexual vs. asexual individuals in response to amputation by analyzing T0 and T3 samples. A DESeq2 model was fitted to gene expression with Time (T0 or T3), reproductive mode (Sexual vs. Asexual) and their interaction as explanatory variables. Genes showing a significant interaction in their expression patterns (implying stronger increase or decrease following amputation in asexual than sexual individuals) were selected based on an adjusted p-value cutoff of <0.05 and |log2FoldChange|>2. To focus on genes of the canonical Wnt/β-catenin pathway, we searched for “canonical wnt” in the Gene Ontology annotations of the transcripts and retained matching transcripts for further inspection. We plotted expression patterns of these selected transcripts based on all time points.

### ALP treatment in amputated animals

Regeneration-deficient sexual animals were used to test the effect of activation of canonical Wnt/β- catenin signalling on regeneration ability. Sexual animals (N=36) 2 weeks after sex induction (lowering temperature from 18 to 8 °C) were head-amputated then randomly divided into two groups: treated with 5 μM alsterpaullone (ALP) or DMSO (the solvent for ALP). ALP treatment lasted for 3 days, after which animals were transferred to fresh hydra medium and development of tentacles was monitored for 14 days. We scored animals regenerated if they developed at least one tentacle on their head, and non-regenerated otherwise. After 14 days, we started feeding animals three times per week and scored their survival 2 months post-amputation.

### ALP treatment and oogenesis

To find out the involvement of canonical Wnt/β-catenin signalling in determining reproductive rate, we treated hydra polyps with ALP at four distinct time points after SI: on the first week (D1 group), second week (D7 group), third week (D14 group) or fourth week (D21 group) after SI (N=6 polyps / week). ALP treatment lasted for three days in each case. Matching controls were treated with DMSO (N=6 polyps / week). Animals were not fed during the treatment, but they received two feedings per week, before and after the treatment. Following treatment, animals were moved to fresh medium, fed three times per week and monitored for egg development (appearance of first egg and total number of detached eggs).

### Statistical analyses

Regeneration and survival rates were analyzed with Fisher’s exact tests. Egg maturation time and fecundity were analyzed with Wilcoxon signed-rank sum tests. P-value adjustments with Benjamini-Hochberg correction were applied to correct for multiple comparisons (i.e. four ALP treatments).

## Results

### Transcriptomic background of regeneration and regeneration loss

After filtering for low quality sequences and contamination from adaptors, rRNA and other species than *H. oligactis*, we obtained a mean ± SD of 18776193 ± 1959587 read pairs (range: 16732138 - 26066578). The assembled transcriptome contained 228212 transcripts; after filtering out low-expression transcripts (<10 count and present in at least 3 samples), this number dropped to 46671. In the Principal Components Analysis sexual and asexual individuals separated clearly from each other (Fig. 1C). In asexual individuals, a shift in gene expression patterns occurred at 4 hpa, which continued at later time points, such that T3 was most different from T0, albeit T1, T2 and T3 samples partly overlapped each other. In sexual individuals, a large shift in gene expression happened at T1, after which samples became similar again to the original expression pattern at T0 (Fig. 1C).

In the DESeq2 analysis, we identified 13 transcripts from the canonical Wnt/β-catenin pathway that showed a significant interaction in their expression patterns between time and reproductive mode: 5 *wnt* genes (*wnt7*, *wnt8, wnt9/10a, wnt9/10c,* and *wnt18*), *wntless* (represented by two transcripts), *axin1*, *dvl2*, *noggin* and *lrrk1* and *p68/c-ets-1* (Fig. 1D). Most *wnt* genes (except *wnt8*), as well as *wntless*, showed reduced expression at T0, but increased substantially at T3 in asexual, but not sexual individuals. For *wnt8*, initial expression was higher in sexual individuals and expression levels declined after amputation in the sexual individuals, while in asexual ones it slightly increased. In the other transcripts (*axin1*, *dvl2*, *noggin* and *lrrk1* and *p68/c-ets-1*), expression levels following amputation also increased more strongly in asexual individuals, while they stayed constant, or even decreased in sexual ones (Fig. 1D). These results show that the canonical Wnt/β-catenin pathway was differently activated in asexual vs. sexual individuals in response to amputation.

### Activation of canonical Wnt/β-catenin signalling restores head regeneration in regeneration-deficient hydra

Given the altered expression pattern of *wnt* genes in sexual and asexual individuals following amputation, we hypothesized that canonical Wnt/β-catenin signalling might be causally involved in the regulation of regeneration loss in *H. oligactis*. In support of that hypothesis, treating sexual polyps with ALP for three days post-amputation significantly increased the proportion of animals that were able to regenerate heads (5.6% of control animals vs 77.8% of treated animals regenerated heads; Fisher’s exact test, p<0.001; Fig. 2B,D). Moreover, after returning to a normal feeding schedule, 72.2% ALP-treated animals recovered and survived, while none of the control animals survived (Fisher’s exact test, p<0.001; Fig. 2C,E). Interestingly, ALP-treated animals that regenerated following amputation and senescence displayed multiple body axes (Fig. 2C).

**Fig. 2.**
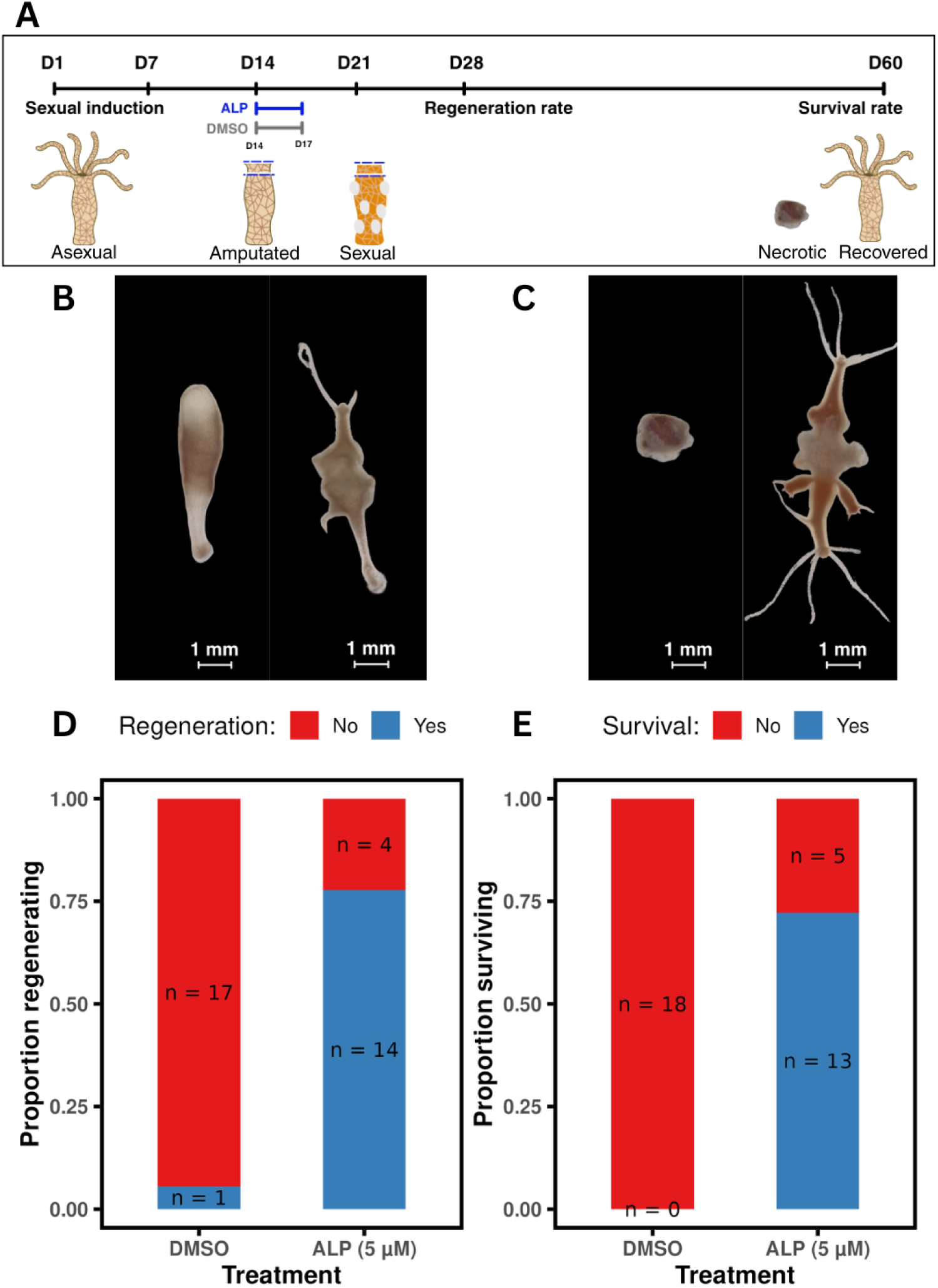
Effects of ALP treatment on head regeneration and subsequent survival. A) Schematic depiction of experimental design. Polyps in early stages of gonadogenesis (two weeks following sexual induction) were amputated below the head and immediately exposed to 5 μM ALP or DMSO for three days. Head regeneration rate was quantified 14 dpa and survival rate quantified 60 days post sexual induction. B) images of control (left) and treated (right) animals at 14 dpa. C) images of control (left) and treated (right) animals 35 dpa. ALP treatment increases head regeneration rate (D) and subsequent survival rate (E).

**Fig. 3.**
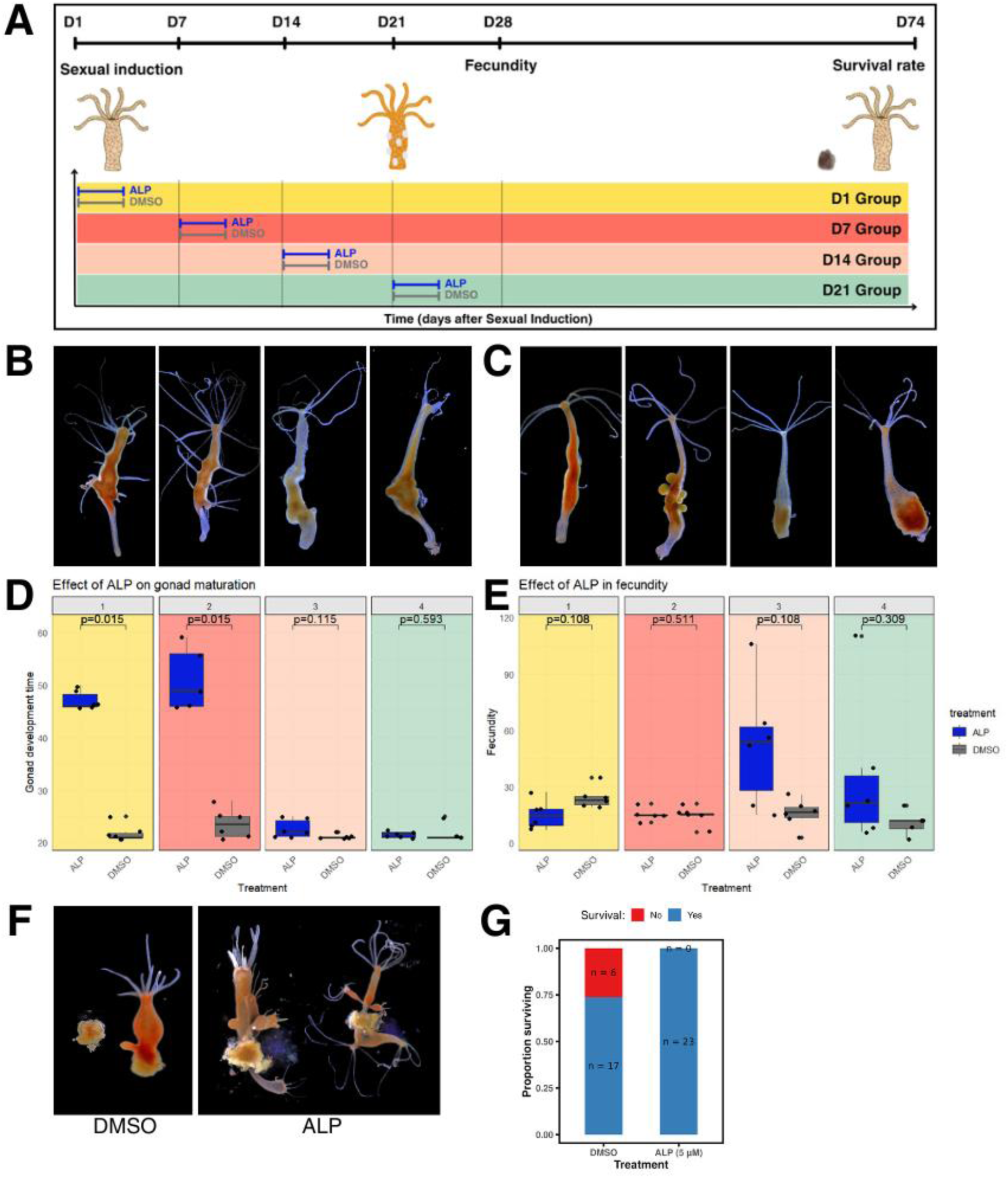
Effects of ALP on egg production and post-reproductive survival. A) Polyps were treated with ALP or DMSO (control) on the first (D1), second (D7), third (D14) or fourth (D21) week after SI, and their egg production and survival monitored until day 75 post-SI. B) Photographs of D1, D7, D14 and D21 ALP-treated polyps (from left to right) two weeks after their treatment. B) Photographs of D1, D7, D14 and D21 control polyps (from left to right) two weeks after their treatment. D) ALP treatment delays the appearance of first eggs in the D1 and D7 groups, but not in the D14 or D21 groups. E) Fecundity is not significantly affected by ALP treatment. F) images of polyps in the control and ALP groups 75 days post-SI. F) ALP treatment increases post- reproductive survival rate.

### Activation of canonical Wnt/β-catenin signalling delays oogenesis and increases survival

Control (DMSO-treated) animals developed eggs 21-28 days after SI, irrespective of when they were treated with DMSO. ALP treated animals, by contrast, showed substantial delay in egg maturation time if treated on the first or the second week post sexual induction (egg maturation time 45-60 days, significantly different from the control groups; Wilcoxon signed rank-sum tests with Benjamini-Hochberg correction, D1 group: p=0.015, D7 group: p=0.015). However, no significant differences were found in the D14 and D21 groups (D14 group: p=0.115, D21 group: p=0.593). These observations suggest that activation of canonical Wnt/β-catenin signalling delays egg production if applied during early stages of gamete development, but later stages do not respond to ALP treatment.

In contrast to egg maturation time, fecundity was not significantly affected by ALP treatment. Animals in the D1 treatment group had slightly fewer eggs than controls, while in groups D3 and D4, treated animals had more eggs than controls. However, none of these differences were statistically significant. Interestingly, we observed that the eggs of D3 and D4 ALP treated animals were smaller than that of controls, suggesting that ALP might prevent oocyte growth through phagocytosis of nurse cells. Overall, these results suggest that activation of canonical Wnt/β- catenin signalling delays egg maturation without having a negative impact on overall fecundity.

All individuals treated with ALP survived until the end of the experiment, but only 74% of the polyps in the control group survived (N=17 out of 23; a significantly lower proportion; Fisher’s exact test, p=0.022).

## Discussion

Canonical Wnt/β-catenin signalling is a central pathway regulating animal development. It is expressed early in embryogenesis, where it determines the development of the oral/aboral body axis, thereby orchestrating embryogenesis^42^, driving adult traits such as asexual reproduction^43^, body size^44^, energy homeostasis^45^, as well as gamete production and fertility^46–48^. In animal species with high rates of regeneration such as sponges, cnidarians, flatworms, acoel worms, fish, amphibians, canonical Wnt/β-catenin signalling is a central determinant of regeneration capacity^18,49–56^. For instance, among planarians, species capable of regeneration have altered expression patterns compared to regeneration-deficient ones and experimental activation of this pathway can restore regeneration ability in species deficient of regeneration^57–59^.

In cnidarians, such as hydra, the role of Wnt signalling in patterning, polarity, cell differentiation and regeneration is well established ^29,60^ Briefly, when cells secrete Wnt, it interacts with specific transmembrane proteins on the surface of other cells (i.e. it has paracrine effects). The Wnt interaction serves as a signal for cells to release β-catenin, a transcription cofactor that translocates to the nucleus to interact with other proteins that together initiate transcription of *wnt* genes. Without Wnt signal, GSK3 phosphorylates β-catenin, leading to its degradation by a proteasome complex^42^. Multiple Wnt ligands are secreted primarily in the apical (head) region of hydra, where they induce a head organizer responsible for the development of oral tissue and tentacles^29,60^. Key Wnt signalling pathway members are activated following injury and drive the regeneration of lost body parts^30,61–63^. Canonical Wnt/β-catenin signalling has been implicated in other developmental processes as well, such as regulating oogenesis^64^. For instance, it has been shown that during oogenesis GSK3 is upregulated in egg-restricted I-cells in *H. vulgaris*. In fact, GSK3 function is required for apoptosis of nurse cells, which is involved in oocyte development ^64^. However, the role of this pathway in determining adult fitness traits in hydra, and its involvement in the regulation of the inducible aging phenotype is unclear. Here we have shown that key Wnt signalling pathway elements are differentially regulated following amputation in asexual and sexual hydra polyps and that experimental activation of canonical Wnt/β-catenin signalling affects multiple components of the inducible aging phenotype, suggesting that shifts in the activity of this pathway during and after gametogenesis is regulating the inducible aging phenotype.

Comparing gene expression profiles of asexual and sexual hydra polyps cultured on the same temperature, we found that asexual polyps starkly upregulated several *wnt* family genes in the tissue below the amputation plane, such as *wnt7*, *wnt9/10a*, *wnt9/10c* and *wnt16*. The upregulation of these transcripts was highest at 72 hpa, much later than in previous studies. However, this is not surprising given that we investigated regeneration at a lower temperature (8 °C), at which cell proliferation and regeneration takes much longer^65,66^. In contrast to the asexual individuals, no upregulation of these genes was observed in the sexual polyps, suggesting that the lack of wnt ligands might contribute to the low rate of regeneration in these animals. A similar pattern was observed in *wntless*, a gene that encodes a protein responsible for transporting *wnt* molecules into the intercellular space. *Dishevelled-2* (*dvl2*) a key component of the transduction of Wnt signals into the cell nucleus was also differentially regulated following amputation in sexual and asexual individuals: its expression decreased with time after amputation in the former and increased in the latter. All these patterns of gene expression suggest reduced activity of Wnt signalling in the sexual individuals.

In support of a *wnt*-mediated loss of regeneration ability in sexual polyps, exposing these individuals to ALP, a *GSK-3* inhibitor that mimics the effects of extracellular Wnt signals restored regeneration ability, as most polyps exposed to ALP were able to regrow tentacles. Some of these polyps also developed ectopic tentacles, as commonly seen in ALP treatments carried out with *H. vulgaris* at higher temperature^67^. The tentacles regenerated on the head were fully functional, and upon receiving food, the animals were able to capture and consume *Artemia*, resulting in high survival rates. In contrast, the controls without tentacles underwent senescence and died. These results clearly demonstrate that a reduced activity of Wnt signalling is driving the regeneration deficiency of sexual polyps.

The loss of regeneration ability of sexual polyps is closely associated with the onset of gonadogenesis^68^. To find out whether Wnt signalling might affect egg production in *H. oligactis,* we treated intact polyps with ALP at various stages of egg development. We found that activation of the canonical Wnt/β-catenin signalling substantially delayed egg production when applied in early stages of oogenesis, while it had little effect later. Oocytes in hydra are produced from a population of germline stem cells derived from multipotent interstitial stem cells that are present in low numbers in asexual polyps cultured at warm temperatures^69,70^. The differentiation of germline stem cells into oocytes happens in several successive stages, which require low temperature^70,71^. During these stages, germline stem cells differentiate into nurse cells, which grow in size and incorporate other nurse cells through phagocytosis, a process inhibited by Wnt signalling^64^. Our observations align with this model of Wnt-dependent egg maturation, as we found that *H. oligactis* females treated with ALP in later stages of oogenesis tended to have smaller eggs. However, we also found that ALP treatment in earlier stages of oogenesis substantially delays egg maturation, the mechanism of which is unclear. First, Wnt activation might have prevented apoptosis of nurse cells and successive growth of oocytes in later stages. However, we find it unlikely that in those early stages apoptosis contributes substantially to oocyte growth. Instead of this, at the earlier stages Wnt signalling might maintain the pluripotency of GSC and inhibit differentiation into gametes, as seen in mouse primordial germ cells^48^. Second, Wnt activation might have slowed down nurse cell and oocyte maturation, resulting in longer egg development type. Third, it is possible that the differentiation pattern of cells was altered, causing stem cells in the early stages of oogenesis to differentiate into cell types other than nurse cells. Supporting this hypothesis, we found that polyps treated with ALP during the early stages of egg maturation developed ectopic tentacles. If some of the cells that would have differentiated into nurse cells in control animals instead differentiated into tentacles, the stem cell pool available for oocyte growth may have been reduced. Further studies are needed to test this hypothesis.

In addition to their altered patterns of oocyte development, sexual polyps with an activated Wnt signalling displayed altered survival patterns following egg production. These polyps did not undergo body size senescence and tissue necrosis like control polyps, retained their feeding capacity, and all of them survived until the end of the experiment. By contrast, polyps in the control group underwent tissue necrosis, shrinkage and lost their feeding ability, resulting in higher mortality. Although the overall difference in survival rates was not very large, the altered pattern of senescence was evident. This relatively low difference in overall survival might have been caused by the large survival rate in the control group. In previous studies with this strain, survival rate in control groups varied from 5 to 40% over the course of several years^68,72,73^. The survival rate observed in this experiment (74%) was much higher. We have several potential explanations to explain these patterns. First, the animals in the current experiment received higher food than in previous experiments (three vs. two feedings per week in previous experiments), which could have enabled them a higher accumulation of resources and increased survival. Alternatively, or additionally, exposure to DMSO and a short starvation period during the treatment might have improved their survival. Indeed, previously we found that reduced food during gametogenesis improved survival in female *H. oligactis*^73^. The role of nutritional status in determining the inducible aging phenotype of *H. oligactis* is an interesting avenue for future research.

In conclusion, activation of canonical Wnt/β-catenin signalling in regeneration-deficient sexual *H. oligactis* polyps restored their regenerative ability, delayed gametogenesis, and improved survival, suggesting that this pathway regulates the inducible aging phenotype in Hydra. The improved survival of polyps with experimentally activated Wnt signalling was not associated with reduced egg production, implying that these polyps were able to maintain both high survival and high fecundity simultaneously. This finding is surprising, as survival and reproduction typically trade off with each other, and longevity mutants often exhibit reduced survival rates, and vice versa (e.g. ^74,75^, but see ^76^). It suggests that hydra polyps can allocate a substantial stem cell pool to reproduction while still retaining enough cells to produce nematocytes for food capture and regenerate their body after massive oocyte production. In this context, the high food levels provided to our experimental animals may have been crucial, as they likely enabled the accumulation of resources for both egg production and survival. Future studies should investigate the mechanisms through which Wnt signalling activation improves survival in hydra.

## Supplementary information

**Supplementary table 1.**
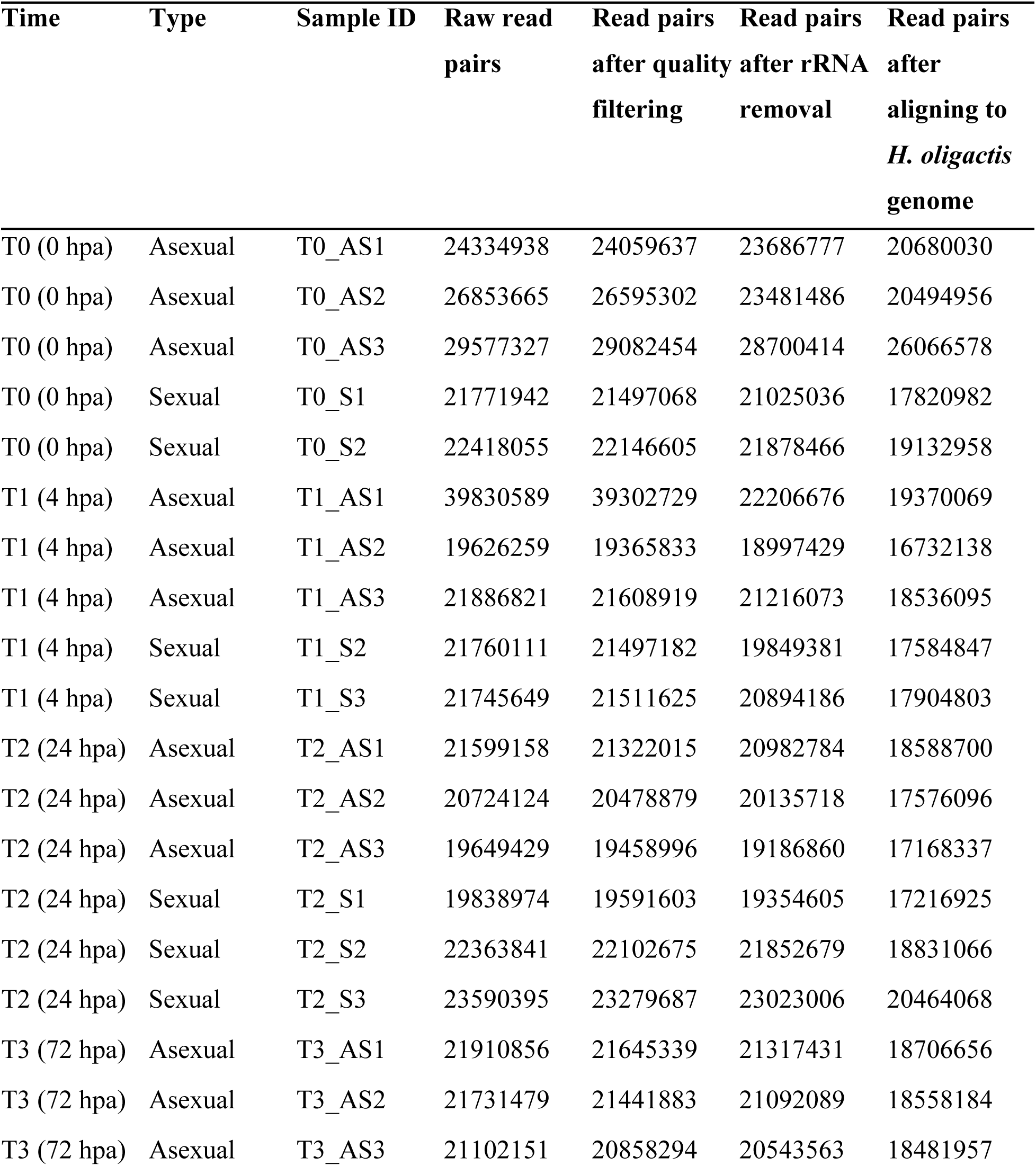
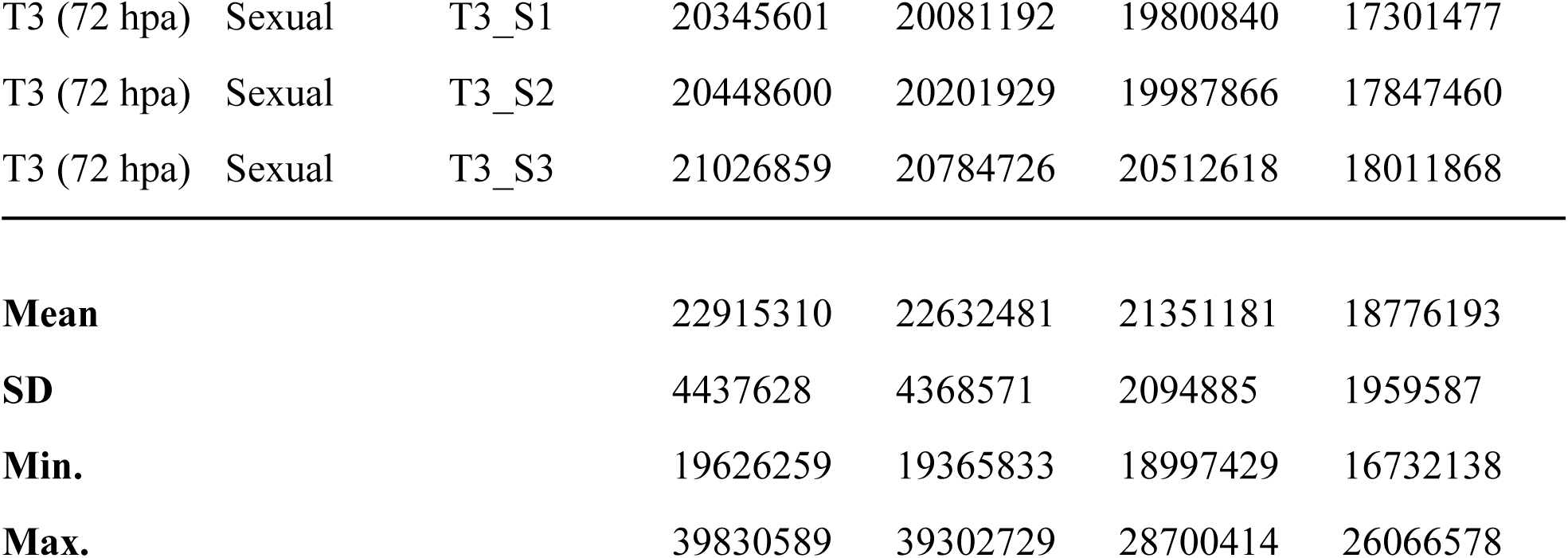
Read statistics for regeneration RNA-Seq time series analysis.

| Time | Type | Sample ID | Raw read<br>pairs | Read pairs<br>after quality<br>filtering | Read pairs<br>after rRNA<br>removal | Read pairs<br>after<br>aligning to<br>*H. oligactis*<br>genome |
| --- | --- | --- | --- | --- | --- | --- |
| T0 (0 hpa) | Asexual | T0_AS1 | 24334938 | 24059637 | 23686777 | 20680030 |
| T0 (0 hpa) | Asexual | T0_AS2 | 26853665 | 26595302 | 23481486 | 20494956 |
| T0 (0 hpa) | Asexual | T0_AS3 | 29577327 | 29082454 | 28700414 | 26066578 |
| T0 (0 hpa) | Sexual | T0_S1 | 21771942 | 21497068 | 21025036 | 17820982 |
| T0 (0 hpa) | Sexual | T0_S2 | 22418055 | 22146605 | 21878466 | 19132958 |
| T1 (4 hpa) | Asexual | T1_AS1 | 39830589 | 39302729 | 22206676 | 19370069 |
| T1 (4 hpa) | Asexual | T1_AS2 | 19626259 | 19365833 | 18997429 | 16732138 |
| T1 (4 hpa) | Asexual | T1_AS3 | 21886821 | 21608919 | 21216073 | 18536095 |
| T1 (4 hpa) | Sexual | T1_S2 | 21760111 | 21497182 | 19849381 | 17584847 |
| T1 (4 hpa) | Sexual | T1_S3 | 21745649 | 21511625 | 20894186 | 17904803 |
| T2 (24 hpa) | Asexual | T2_AS1 | 21599158 | 21322015 | 20982784 | 18588700 |
| T2 (24 hpa) | Asexual | T2_AS2 | 20724124 | 20478879 | 20135718 | 17576096 |
| T2 (24 hpa) | Asexual | T2_AS3 | 19649429 | 19458996 | 19186860 | 17168337 |
| T2 (24 hpa) | Sexual | T2_S1 | 19838974 | 19591603 | 19354605 | 17216925 |
| T2 (24 hpa) | Sexual | T2_S2 | 22363841 | 22102675 | 21852679 | 18831066 |
| T2 (24 hpa) | Sexual | T2_S3 | 23590395 | 23279687 | 23023006 | 20464068 |
| T3 (72 hpa) | Asexual | T3_AS1 | 21910856 | 21645339 | 21317431 | 18706656 |
| T3 (72 hpa) | Asexual | T3_AS2 | 21731479 | 21441883 | 21092089 | 18558184 |
| T3 (72 hpa) | Asexual | T3_AS3 | 21102151 | 20858294 | 20543563 | 18481957 |
| T3 (72 hpa) | Sexual | T3_S1 | 20345601 | 20081192 | 19800840 | 17301477 |
| T3 (72 hpa) | Sexual | T3_S2 | 20448600 | 20201929 | 19987866 | 17847460 |
| T3 (72 hpa) | Sexual | T3_S3 | 21026859 | 20784726 | 20512618 | 18011868 |
| **Mean** |  |  | 22915310 | 22632481 | 21351181 | 18776193 |
| **SD** |  |  | 4437628 | 4368571 | 2094885 | 1959587 |
| **Min.** |  |  | 19626259 | 19365833 | 18997429 | 16732138 |
| **Max.** |  |  | 39830589 | 39302729 | 28700414 | 26066578 |

